# Probing the proton release by Photosystem II in the S_1_ to S_2_ high-spin transition

**DOI:** 10.1101/2022.01.19.476895

**Authors:** Alain Boussac, Miwa Sugiura, Julien Sellés

**Author notes:** **Abbreviations:** Chl, chlorophyll; Chl_D1_/Chl_D2_, accessory Chl’s on the D1 or D2 side, respectively; PSII, Photosystem II; MES, 2–(*N*–morpholino) ethanesulfonic acid; HEPES, 4-(2-hydroxyethyl)-1-piperazine ethane sulfonic acid. P_680_, primary electron donor; P_D1_ and P_D2_; Chl monomer of P_680_ on the D1 or D2 side, respectively, Phe_D1_ and Phe_D2_, pheophytin on the D1 or D2 side, respectively; Q_A_, primary quinone acceptor; Q_B_, secondary quinone acceptor; Tyr_Z_, redox active tyrosine 161 of D1; WT*3, *T. elongatus* mutant strain containing only the *psbA_3_* gene and a His_6_-tag on the C-terminus of CP43. EPR, Electron Paramagnetic Resonance spectroscopy; EDNMR, ELDOR-detected Nuclear Magnetic Resonance; ELDOR, Electron-Electron Double Resonance; SQR, sum of the squares of the residues.

## Abstract

The stoichiometry and kinetics of the proton release were investigated during each transition of the S-state cycle in Photosystem II (PSII) from *Thermosynechococcus elongatus* containing either a Mn_4_CaO_5_ (PSII/Ca) or a Mn_4_SrO_5_ (PSII/Sr) cluster. The measurements were done at pH 6.0 and pH 7.0 knowing that, in PSII/Ca at pH 6.0 and pH 7.0 and in PSII/Sr at pH 6.0, the flash-induced S_2_-state is in a low-spin configuration (S_2_^LS^) whereas in PSII/Sr at pH 7.0, the S_2_-state is in a high-spin configuration (S_2_^HS^) in half of the centers. Two measurements were done; the time-resolved flash dependent *i*) absorption of either bromocresol purple at pH 6.0 or neutral red at pH 7.0 and *ii*) electrochromism in the Soret band of P_D1_ at 440 nm. The fittings of the oscillations with a period of four indicate that one proton is released in the S_1_ to S_2_^HS^ transition in PSII/Sr at pH 7.0. It has previously been suggested that the proton released in the S_2_^LS^ to S_3_ transition would be released in a S_2_^LS^Tyr_Z_^●^ → S_2_^HS^Tyr_Z_^●^ transition before the electron transfer from the cluster to Tyr_Z_^●^ occurs. The release of a proton in the S_1_Tyr_Z_^●^ →S_2_^HS^Tyr_Z_ transition would logically imply that this proton release is missing in the S_2_^HS^Tyr_Z_^●^ to S_3_Tyr_Z_ transition. Instead, the proton release in the S_1_ to S_2_^HS^ transition in PSII/Sr at pH 7.0 was mainly done at the expense of the proton release in the S_3_ to S_0_ and S_0_ to S_1_ transitions. However, at pH 7.0, the electrochromism of P_D1_ seems larger in PSII/Sr when compared to PSII/Ca in the S_3_ state. This points to the complex link between proton movements in and immediately around the Mn_4_ cluster and the mechanism leading to the release of protons into the bulk.

## Introduction

Oxygenic photosynthesis is responsible for most of the energy input on Earth by converting the solar energy into fibers, foods and fuels. This process occurs in cyanobacteria, algae and plants. Photosystem II (PSII), the water-splitting enzyme, is at the heart of this process; see for example [1] for a recent view with an evolutionary perspective.

Mature PSII generally consists of 20 subunits with 17 trans-membrane and 3 extrinsic membrane proteins. The PSII binds 35 chlorophylls *a* (Chl-*a*), 2 pheophytins (Phe), 1 membrane b-type cytochrome, 1extrinsic c-type cytochrome (in cyanobacteria and red algae), 1 non-heme iron, 2 plastoquinones (Q_A_ and Q_B_), the Mn_4_CaO_5_ cluster, 2 Cl^-^, 12 carotenoids and 25 lipids [2, 3]. Recently, the 4^th^ extrinsic PsbQ subunit has also been found in PSII from *Synechocystis* sp. PCC 6803 in addition to PsbV, PsbO and PsbU [4].

Among the 35 Chls, 31 are antenna Chls and 4 (P_D1_, P_D2_, Chl_D1_ and Chl_D2_) together with the 2 Phe molecules constitute the reaction center of PSII. After the absorption of a photon by the antenna, the excitation energy is transferred to the photochemical trap that consists of the four Chls; P_D1_, P_D2_, Chl_D1_, Chl_D2_. After a few picoseconds, a charge separation occurs resulting ultimately in the formation of the Chl_D1_^+^Phe_D1_^-^ and then P_D1_^+^Phe_D1_^-^ radical pair states [5, 6].

After the charge separation, P_D1_^+^ oxidizes Tyr_Z_, the Tyr161 of the D1 polypeptide, which then is reduced by the Mn_4_CaO_5_ cluster, see [7] for one recent review. The electron on Phe_D1_^-^ is then transferred to Q_A_, the primary quinone electron acceptor, and then to Q_B_, the second quinone electron acceptor. Whereas Q_A_ can be only singly reduced under normal conditions, Q_B_ accepts two electrons and two protons before to leave its binding site and to be replaced by an oxidized Q_B_ molecule from the membrane plastoquinone pool, [8–11] and references therein.

The Mn_4_CaO_5_ cluster, oxidized by the Tyr_Z_^●^ radical after each charge separation, cycles through five redox states denoted S*_n_*, where *n* stands for the number of stored oxidizing equivalents. The S_1_-state is stable in the dark and therefore S_1_ is the preponderant state upon dark-adaptation. When the S_4_-state is formed after the 3^rd^ flash of light given on dark-adapted PSII, two water molecules bound to the cluster are oxidized, O_2_ is released and the S_0_-state is reformed, [12, 13].

Thanks to the advent of serial femtosecond X-ray free electron laser crystallography, structures of the Mn_4_CaO_5_ cluster have been resolved in the dark-adapted state with the 4 Mn ions in a redox state as close as possible to that in the S_1_ state, *i.e.* Mn^III^_2_Mn^IV^ [3, 14]. The Mn_4_CaO_5_ structure resemble a distorted chair including a μ-oxo-bridged cuboidal Mn_3_O_4_Ca unit with a fourth Mn attached to this core structure *via* two μ-oxo bridges involving the two oxygen’s O4 and O5. After further refinement of the electron density maps, some works could indicate an heterogeneity in the valence of Mn ions between the two PSII monomers and/or could show a lower than expected oxidation state of some Mn ions [15–17]. However, other groups [18–20] do not share such a conclusion mainly by taking into account the Jahn–Teller effect on the orientation of the bondings of the Mn^III^.

Recently, important progresses have been done in the resolution of the crystal structures in the S_2_ and S_3_ states [21–24]. The structural changes identified in the S_1_ to S_2_ transition and to a greater extent in the S_2_ to S_3_ transition are too numerous to detail them in a few lines. Very briefly, these works show that, in S_2_, the changes in the structure of the cluster are minor and more or less correspond to those expected for the valence change of the Mn4 from +III to +IV in the context of the high-valence option [18]. Importantly, water molecules in the “O1” and “O4” channels, defined as such because they start from the O1 and O4 oxygens of the cluster, appeared localized slightly differently in S_2_ than in S_1_. In contrast, in the S_2_ to S_3_ transition, major structural changes have been detected together with the insertion of a 6^th^ oxygen (O6 or Ox), possibly W3 originally bound to the Ca site, bridging Mn1 and Ca. This oxygen is supposed to correspond to the second water substrate molecule and is close to the bridging oxygen O5 supposed to be the first water substrate molecule [25]. An important movement of the Glu189 residue would allow its carboxylate chain to make a hydrogen bond with the protonated form of this 6^th^ oxygen in S_3_ [21].

EPR data studies, supported by computational works, show the existence of more than one structural forms of S_1_, S_2_ and S_3_. The two S_1_ EPR signals seen with a parallel mode detection at *g* ∼ 4.8 and *g* ∼ 12 [26–28] were recently attributed to an orientational Jahn–Teller isomerism of the dangler Mn4 with the valence +III [20]. The authors in [20] further suggested that this isomerism in the S_1_ state is at the origin of the valence isomerism in the S_2_-state. Indeed, depending on the conditions, at least two S_2_ EPR signals can be detected at helium temperatures. The first one has a low-spin S  =  1/2 value, S_2_^LS^, characterized by a multiline signal made up of at least 20 lines separated by approximately 80 gauss, centered at *g* ∼ 2.0 and spread over roughly 1800 gauss [29–31]. The second configuration of S_2_ is a high-spin ground state, S_2_^HS^, with *S*  ≥  5/2. In plant PSII, S_2_^HS^ may exhibit either a derivative-like EPR signal centered at *g*∼ 4.1 [32, 33] or more complex signals at higher *g* values [34, 35]. In cyanobacterial PSII, the S_2_^HS^ EPR signal has a derivative-like EPR signal centered at *g*∼ 4.8 [36, 37].

In a computational work [38], it was proposed that the *g* ∼ 4.1 signal in plant PSII have almost the same coordination and environment as the *g* ∼ 2.0 signal but with the Mn^III^ ion located on the dangler Mn4 in the S_2_^HS^ state instead on the Mn1 in the S_2_^LS^ state. This valence swap would be accompanied by a moving of the oxygen O5 from a position where it links the Mn4, Mn3 and the Ca in the S_2_^LS^ configuration to a position where it bridges the Mn1, Mn3 and Ca ions in the S_2_^HS^ configuration resulting into the so called closed cubane structure. In other computational works [39, 40] the authors arrived at a very different model in which, starting from the S_2_^LS^ configuration, the protonation of O4 would lead to an *S* = 5/2 ground state when W1, one of the two water molecules with W2 bound to the dangler Mn4, is present as an aquo ligand. The further deprotonation of W1 to form a hydroxo ligand would then give rise to an *S* = 7/2 ground state [39, 40]. In the same work, it was proposed that the form *S* = 7/2 was probably that needed to progress to S_3_. The fact that the closed cubane S_2_ structure defined in [41, 42] is not necessary to progress to S_3_ is in line with some earlier works [43]. Experimentally [36, 37], the S_2_^HS^ form able to progress to S_3_ at low temperatures is the *g* ∼ 4.8 form, *i.e.* the *S* = 7/2 form that corresponds to an open cubane structure in [39, 40].

From an EPR point of view, the S_3_-state also exhibits some heterogeneities. The majority of centers exhibit a spin *S* = 3 ground state [44–46]. In this *S* = 3 configuration, the four Mn ions of the cluster have a Mn^IV^ formal oxidation state with an octahedral ligation sphere in an open cubane structure [46]. In this model, the dangler Mn^IV^ (*S* = 3/2) is antiferromagnetically coupled to the open cubane motif (Mn^IV^)_3_ with *S =* 9/2. The remaining centers are EPR invisible, *e.g.* [37]. The relationship between the different S_2_ and S_3_ configurations was recently investigated by EPR spectroscopy [37] and discussed in the context of the structural heterogeneities proposed from computational works [41]. In [41], the high-spin S_2_-state was considered to be a closed cubane structure something that needs to be reconsidered in view of the proposals made in [39, 40] and because there is less and less structural supports for the closed cubane structure, *e.g.* [22].

A third S_3_ configuration with a broadened S_3_ signal was identified with EDNMR in the presence of glycerol [47, 48] and in PSII/Sr [49]. Although in [47] the authors did not completely rule out the presence of a closed cubane, five-coordinate S_3_ form, at the origin of this EPR signal, they favored a perturbation of the coordination environment at Mn4 and/or Mn3 in an open cubane S_3_ structure induced by glycerol. With X-and Q-band EPR experiments performed in the S_3_-state of plant PSII, both in the perpendicular and parallel modes, a high-spin, *S =* 6, was shown to coexist with the *S =* 3 configuration. This *S =* 6 form was attributed to a form of S_3_ without O6/Ox bound and with the Mn^IV^ part of the cluster in ferromagnetic interaction with the unsaturated dangler Mn^IV^ [48]. Nevertheless these two forms of S_3_ are not detected by X-band EPR, it seems unlikely that they correspond to the EPR invisible S_3_ mentioned above. Indeed, the new S_3_ signals described in [47, 48] are detectable in the presence of glycerol and methanol, whereas the formation of the (S_2_Tyr_Z_^●^)’ state upon a near-IR illumination in the centers in S_3_ defined as EPR invisible is inhibited in the presence of glycerol (and in the presence of methanol in plant PSII) [50, 51]. In cyanobacterial PSII/Sr [49], a proportion of centers exhibited a pulsed W-band field-swept S_3_ spectrum much broader than in PSII/Ca. This signal was proposed to be present in centers containing a 5-coordinate Mn ion in centers in which no water binding event takes place during the S_2_ to S_3_ transition. It was therefore proposed that, in these centers, the oxidation event would precede the water binding. Computational works also suggested heterogeneities in S_3_ [52–54] with also a *S =* 6 spin value [54, 55].

None of the heterogeneities described above were detected in the crystallographic structures of S_2_ and S_3_ known to date [21–24]. It is quite possible that some of the structural differences that cause the differences identified in EPR are too small to be detectable given the resolution of the crystallographic data. This at least shows, if it were necessary, that spectroscopy remains an indispensable complement to crystallography.

The EPR data summarized above describes a static view of the trapped configurations. Kinetically, it is well documented that the transition from S_2_ to S_3_ involved at least two phases. The fastest phase, with a *t*_1/2_ ≤ 25 µs, is attributed to a proton transfer/release. This fast phase precedes the electron transfer from S_2_ to the Tyr_Z_^●^, which occurs with a *t*_1/2_ ≤ 300 µs [56–60], and the binding of O6/Ox to the Mn1, *e.g.* [21]. It was discussed earlier that the fast phase could correspond to the release of a proton in an intermediate step S_2_^LS^Tyr_Z_^●^→ S_2_^HS^Tyr_Z_^●^ before the S_2_^HS^Tyr_Z_^●^ → S_3_*^S^*^=3^ transition occurs [36]. The existence of intermediate states in the S_2_ to S_3_ transition was tracked by following, at room temperature, the structural changes in the S_2_ to S_3_ transition at time points from 50 μs to 200 ms after the 2^nd^ flash [21]. Although no indication was found for a closed cubane intermediate this says, for the moment, nothing on the spin state of the intermediate forms of S_2_ able to progress to S_3_. Indeed, we have seen above that the S_2_^HS^ may have an open cubane structure, *e.g.* [39, 40]. In addition, a transient state, by definition, has a low concentration that makes its detection difficult. In an even more recent work [22], a structural dynamics in the water and proton channels was highlighted during the S_2_ to S_3_ transition.

The question of the existence of a high spin intermediate state in the S_2_ to S_3_ transition remains. In particular, it is questionable whether the fast phase observed in this transition corresponds to a proton release/movement associated with the formation of a S_2_^HS^ state, as has been suggested [36, 60]. If this is correct, we would expect to detect a change in the flash pattern of the proton release in conditions in which the S_2_ *g*∼ 4.8 EPR signal is the flash-induced state. In order to probe this model, we have kinetically followed the flash dependent proton release in PSII/Ca and PSII/Sr at pH 6.0 and 7.0, knowing that at pH 7.0 half of the centers exhibit the S_2_^HS^ signal at *g*∼ 4.8 in PSII/Sr in contrast to PSII/Ca. This was done by recording absorption changes of pH-responding dyes, *i.e.* bromocresol purple at pH 6.0 and neutral red at pH 7.0, *e.g.* [61, 62]. As a control experiment, changes in the electrostatic environment of P_D1_ undergone upon the reduction/oxidation and deprotonation/protonation reactions of either the Mn_4_CaO_5_ cluster or the Mn_4_SrO_5_ cluster were also followed by recording the time-resolved absorption change differences 440 nm-*minus*-424 nm during the S-state cycle, *e.g.* [60, 63].

## Materials and Methods

The *Thermosynechococcus elongatus* strain used was the *ΔpsbA_1_*, *ΔpsbA_2_* deletion mutant, referred to as either WT*3-PSII or PsbA3-PSII [64]. This strain was constructed from the *T. elongatus* 43-H strain that had a His_6_-tag on the carboxy terminus of CP43 [65]. The cells were cultivated in the presence of either Ca^2+^ or Sr^2+^ [66]. PSII/Ca and PSII/Sr purifications were achieved with a protocol previously described [67].

Absorption changes measurements were done with a lab-built spectrophotometer [68] slightly modified as previously described [67]. The 440 nm-*minus*-424 nm experiments were performed as previously reported [60]. For that, the samples were diluted in 1 M betaine, 15 mM CaCl_2_, 15 mM MgCl_2_, and either 40 mM Mes at pH 6.0 or 40 mM Hepes at pH 7.0. The pH of the two solutions was adjusted with NaOH. The measurements with the dyes were done as previously reported [67, 69]. In this case, the samples were diluted in 1 M betaine, 15 mM CaCl_2_, 15 mM MgCl_2_, and either 150 µM bromocresol purple, *i.e.* 312 times the concentration of PSII, at pH 6.0 or 40 µM neutral red at pH 7.0, *i.e.* 50 times the concentration of PSII. PSII samples were dark-adapted for ∼ 1 h at room temperature (20–22°C) before the addition of 100 µM phenyl *p*–benzoquinone (PPBQ) dissolved in dimethyl sulfoxide and 100 µM ferricyanide. The pH of the stock solutions of ferricyanide (100 mM) was adjusted to either pH 6.0 or pH 7.0 prior to their addition to the samples. The chlorophyll concentration of all the samples was ∼ 25 µg of Chl/mL. After the ΔI/I measurements, the absorption of each diluted batch of samples was precisely controlled to avoid errors due to the dilution of concentrated samples. The ΔI/I values were then normalized to *A*_673_ = 1.75, that is very close to 25 µg Chl/mL with ε ∼ 70 mM^- 1^·cm^-1^ at 674 nm for dimeric PSII [70].

The fittings of the data were done as previously described [60,63,66,71] with the additional details given in the text by using the Excel solver. All the data are the average of 2 to 3 experiments done on different batches of PSII resulting from different purifications. The noise defined as the maximum variation of the averaged signal before the first actinic flash was ∼ ± 70 10^-6^ ΔI/I units at 575 nm, ∼ ± 30 10^-6^ ΔI/I units at 547 nm and ∼ ± 20 10^-6^ ΔI/I units for the 440 nm-*minus*-424 nm difference.

## Results

### Stoichiometry of the S-state dependent proton release

After one-flash illumination given at pH 6.0 to dark-adapted PSII, *i.e.* in the S_1_-state, both PSII/Ca and PSII/Sr exhibit a *S =* 1/2 low-spin configuration, S_2_^LS^ [36] At pH 7.0, the S_2_^LS^ state is the flash-induced state in PSII/Ca whereas in PSII/Sr a proportion of centers exhibits a S_2_-state in a *S* ≥ 5/2 high-spin configuration, S_2_^HS^. In our previous work [36], in which the PSII was first washed in a buffer-free medium and then the pH was directly adjusted in the EPR tubes by adding 100 mM of different buffers, we found that the pK value of the S_2_^LS^ ↔ S_2_^HS^ equilibrium was ∼ 7.5. However, after this earlier work, we have found that such a protocol overestimated the pH values by ∼ 0.5 pH unit so that the pK of the S_2_^LS^ ↔ S_2_^HS^ equilibrium is actually ∼ 7.0 instead of 7.5. The data at pH 7.0 in PSII/Sr in this work therefore correspond to a situation in which 50 % of the centers are in S_2_ ^HS^ state (and this has been controlled by EPR with PSII/Sr washed with a large volume of the medium at pH 7.0, not shown). It would have been much more comfortable for the interpretation of the results to do the experiment at pH values where the proportion of S_2_^HS^ is greater. Unfortunately, for higher pH values than 7.0, the integrity of the PSII/Sr estimated by recording the amplitude of the S_2_^LS^ and S_2_^HS^ EPR signals after variable times (not shown), and by following the amplitude of the period four oscillations (not shown), decreases much faster than the time required for doing the experiments reported here (about 2 hours and half). This problem did not occurred in previous EPR experiments [36] since in this case the samples were frozen immediately after the addition of the buffers.

Fig. 1, shows the absorbance changes of either bromocresol purple at pH 6.0 (purple full circles in Panels A and B) or neutral red at pH 7.0 (red full circles in Panels C and D). The ΔI/I were measured 100 ms after each flash, 400 ms apart, of the series. The measurements were done with PSII/Ca in Panels A and C and with PSII/Sr in Panels B and D. A proton released by PSII, *i.e.* an acidification of the bulk, results in a decrease of the absorption at 575 nm with bromocresol purple and in an increase of the absorption at 547 nm with neutral red. The wavelength of 547 nm was chosen because it corresponds to the isosbestic point of the electrochromic blue shift undergone by Phe_D1_ in the Q_X_ absorption region upon the formation of Q_A_^-^ with PsbA3 as the D1 protein, *e.g.* [60].

**Figure 1:**
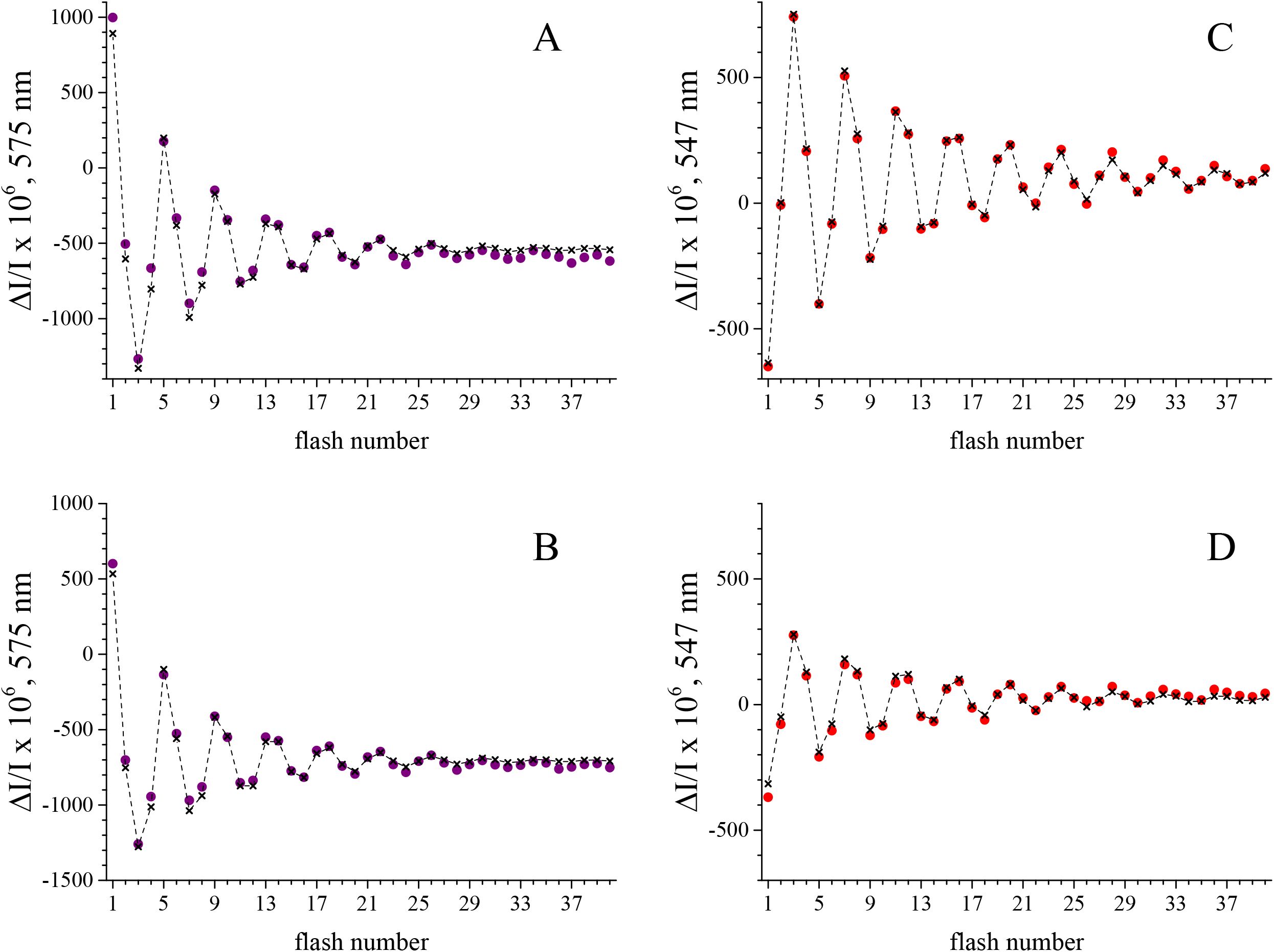
Sequence of the amplitude of the absorption changes using a series of saturating flashes (spaced 400 ms apart). The full circles, shows the absorbance changes at 575 nm of either bromocresol purple (150 µM) at pH 6.0 (Panels A and B, purple full circles) or at 547 nm of neutral red (40 µM) at pH 7.0 (Panels C and D, red full circles). The measurements were done with PSII/Ca in Panels A and C and with PSII/Sr in Panels B and D. The samples ([Chl] = 25 µg mL^-1^) were dark-adapted for 1 h at room temperature before the addition of 100 µM PPBQ dissolved in dimethyl sulfoxide and 100 µM ferricyanide. The ΔI/I were measured 100 ms after each flash. The black crosses joined by a continuous line are the fits of the data with the parameters listed in Table 1.

**Table 1:**
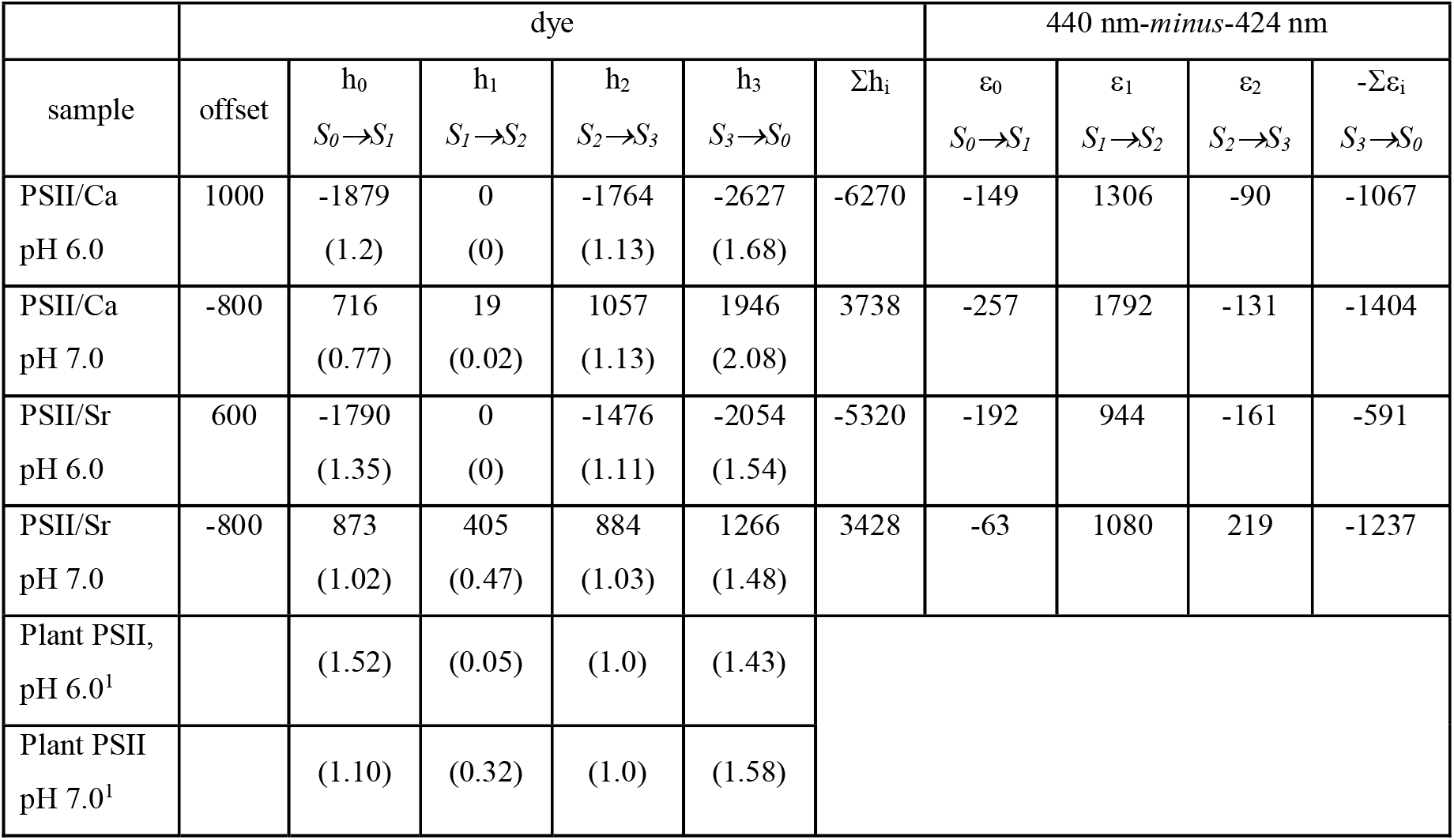
Fitted values for the data shown in Figures 1 and 3. ΔI/I x 10^6^ values corresponding to the best fittings shown in Figures 1 and 3. The h_0_, h_1_, h_2_ and h_3_ values are the ΔI/I changes associated to the changes in the protonation state of the dyes in the S_0_ to S_1_, S_1_ to S_2_, S_2_ to S_3_ and S_3_ to S_0_ transitions, respectively. The Σh_i_ value corresponds to the total absorption change for one turnover (*i.e.* h_0_ + h_1_ + h_2_ + h_3_) which corresponds to 4 H^+^. The values in brackets are the ratio h_i_/Σh_i_ which corresponds to stoichiometry of the proton released in the corresponding transition. The ε_0_, ε_1_, ε_2_ values are the extinction coefficients corresponding to the S_0_ to S_1_, S_1_ to S_2_ and S_2_ to S_3_ transitions, respectively. By definition [63], the extinction coefficient of the S_3_ to S_0_ transition is -Σε_I_ (*i.e.* –(ε_0_ + ε_1_ + ε_2_)). In all the fittings the miss parameter, considered as identical in all the transitions, was found to be between 8 and 10 % and the proportion of centers in S_1_ between 90 and 100 %. ^1^Data taken in [62].

In PSII/Ca, at pH 6.0 (Panel A), there was a large increase in the absorption after the first flash. Since no proton release is supposed to occur in the S_1_ to S_2_ transition in such a sample at this pH, this change in the ΔI/I mainly corresponds to a proton uptake coupled to the reduction of the oxidized non-heme iron by Q_A_ ^-^. After the 5^th^ flash, the ΔI/I change remained positive. This clearly indicates that there was also a proton uptake after this 5^th^ flash (mainly the S_1_ to S_2_ transition during the second turnover). In PSII/Ca at pH 7.0 (Panel C), with neutral red, this absorption due to the proton uptake coupled to the reduction of the oxidized non-heme iron seemed much more pronounced. Indeed, at least until the 21^st^ flash there was a negative contribution of the ΔI/I after the 3^rd^ and 4^th^ flashes, the 7^th^ and 8^th^, and so on, flashes. In the analysis of such experiments, the data from the first flash are usually discarded due to several non-oscillating contributions [63]. In previous works, we did not consider that the non-heme iron could contribute after the first flash [67,69,71]. The data in Panel A and C in Fig. 1 show that this is not completely true under the present conditions. Although the conclusions made previously remain valid, the smallness of the effects that are expected here require to take into account the proton uptake coupled to the reduction of the non-heme iron after each flash of the series. To this end, the stoichiometry of the proton release was therefore estimated for different amounts of proton uptake occurring on each flash, assuming that this proton uptake is the same on all flashes from the 2^nd^ onwards. This procedure is more or less equivalent to adding a constant offset to the proton release calculated as described previously [71]. This offset also takes into account the slow drift that was equally present on all flashes. This drift has an unknown origin and may vary from experiment to experiment. Part of this process probably involves the release of a proton due to the reoxidation of non-heme iron during the dark period between flashes.

Fig. 2 shows the results of these fittings at pH 6.0 for PSII/Ca in Panel A and for PSII/Sr in Panel B. Panel C and D shows the fittings at pH 7.0 for PSII/Ca and PSII/Sr, respectively. The X-axis corresponds to the different values of the offset, in ΔI/I units, which were tested in the fitting procedure. This offset corresponds to the sum of the proton uptake following the reduction of the non-heme iron and the drift 100 ms after the flash. The values h_0_ (green curve), h_1_ (black curve), h_2_ (red curve), h_3_ (blue curve) are the fitted ΔI/I corresponding to the release of proton(s) in the S_0_ to S_1_, S_1_ to S_2_, S_2_ to S_3_ and S_3_ to S_0_ transitions, respectively, for each offset value. The yellow curves correspond to the sum of the squares of the residues (SQR) calculated from the 2^nd^ to the 40^th^ flash. At pH 6.0, the SQR was multiplied by -1 for a better visualisation of the curves in the graph.

**Figure 2:**
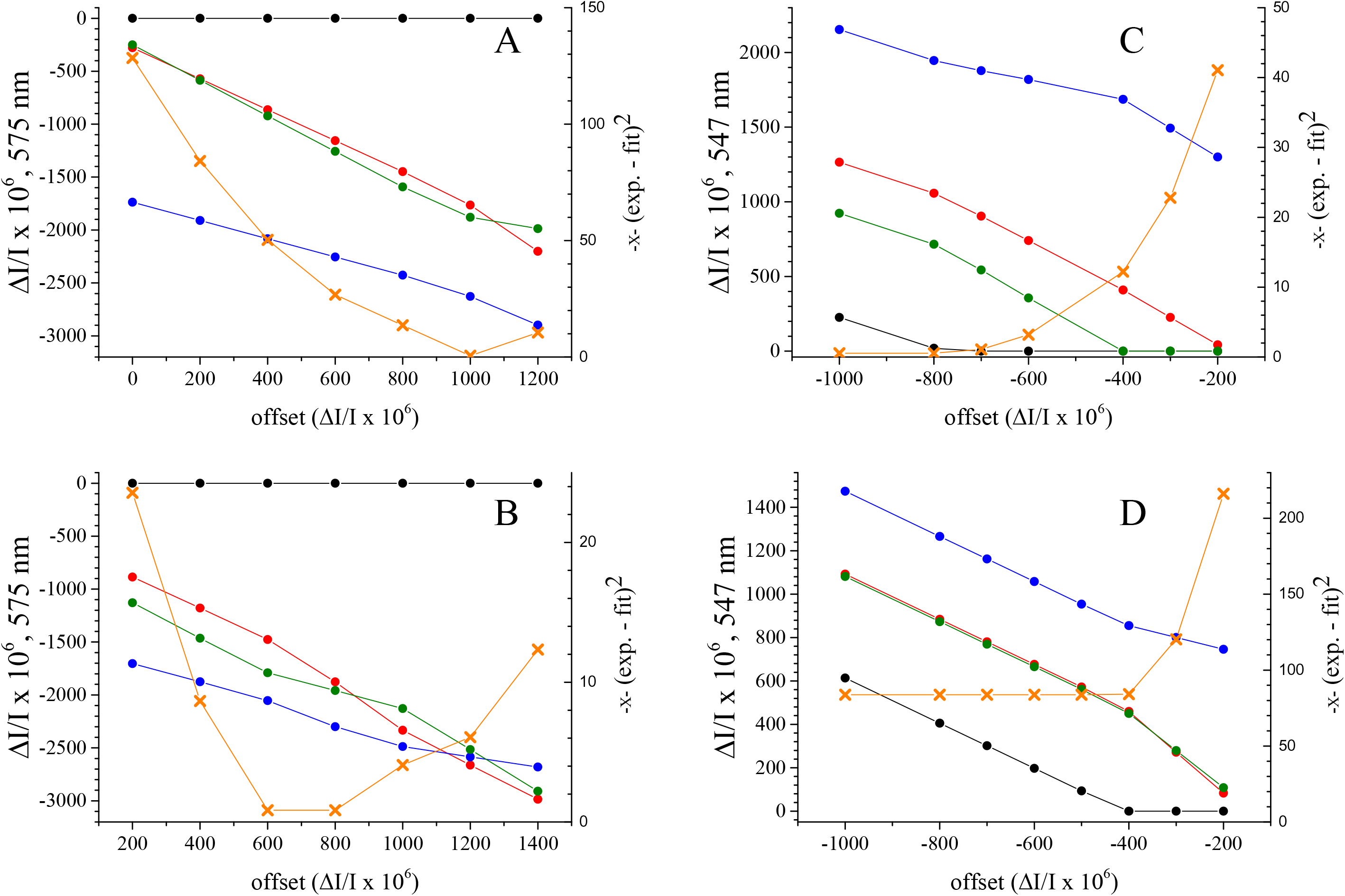
Results of the fittings at pH 6.0 for PSII/Ca in Panel A and for PSII/Sr in Panel B and at pH 7.0 for PSII/Ca in Panel C and for PSII/Sr in Panel D. The X-axis corresponds to different values of the offset, in ΔI/I units, which were tested in the fitting procedure. This offset corresponds to the sum of the proton uptake and the amplitude of the drift 100 ms after the flash. The values h_0_ (in green), h_1_ (in black), h_2_ (in red), h3 (in blue) are the fitted ΔI/I values corresponding to the release of proton(s) in the S_0_ to S_1_, S_1_ to S_2_, S_2_ to S_3_ and S_3_ to S_0_ transitions, respectively, for each value of the offset. The yellow curve corresponds to the sum of the squares of the residues calculated from the 2^nd^ to the 40^th^ flash. At pH 6.0, the sum of the squares of the residues was multiplied by -1 for a better visualisation of the curves in the graph.

At pH 6.0, in both PSII/Ca and PSII/Sr, whatever the offset value, there was no proton release in the S_1_ to S_2_ transition (black curves). For offset values resulting in the smallest SQR, the proton release estimated from the relative amplitudes of the ΔI/I was similar in the S_2_ to S_3_ (red curves) and S_0_ to S_1_ (green curves) transitions and almost twice in the S_3_ to S_0_ transition (blue curves) as generally found for the pattern of the proton release [61, 62].

At pH 7.0, in PSII/Ca, when the SQR was the smallest, the proton release on the S_1_ to S_2_ transition remained either zero (or very small if not zero). In the S_0_ to S_1_ transition, the proton release became smaller than in the S_2_ to S_3_ transition as observed in PSII from plant [62]. In PSII/Sr, at pH 7.0, the proton release in the S_1_ to S_2_ transition significantly increased to ∼ 0.5 and that constitutes the main result of this experiment. These observations will be further discussed with the analysis of the kinetics in Fig. 4. However, we can already note that the larger proton release in the S_1_ to S_2_ transition in PSII/Sr at pH 7.0 was done in part at the expense of the proton release in the S_3_ to S_0_ and S_0_ to S_1_ transitions when compared to the situation in PSII/Ca at pH 7.0. Table 1 summarizes the results of the fittings. The crosses joined by dashed lines in Fig. 1 are the fits using the values reported in Table 1. The sums of the ΔI/I for an offset of ∼ -800 at pH 7.0 are comparable in PSII/Ca and PSII/Sr which shows that the apparent smaller amplitudes of the oscillations at this pH are due to a different stoichiometry of the proton release.

**Figure 3:**
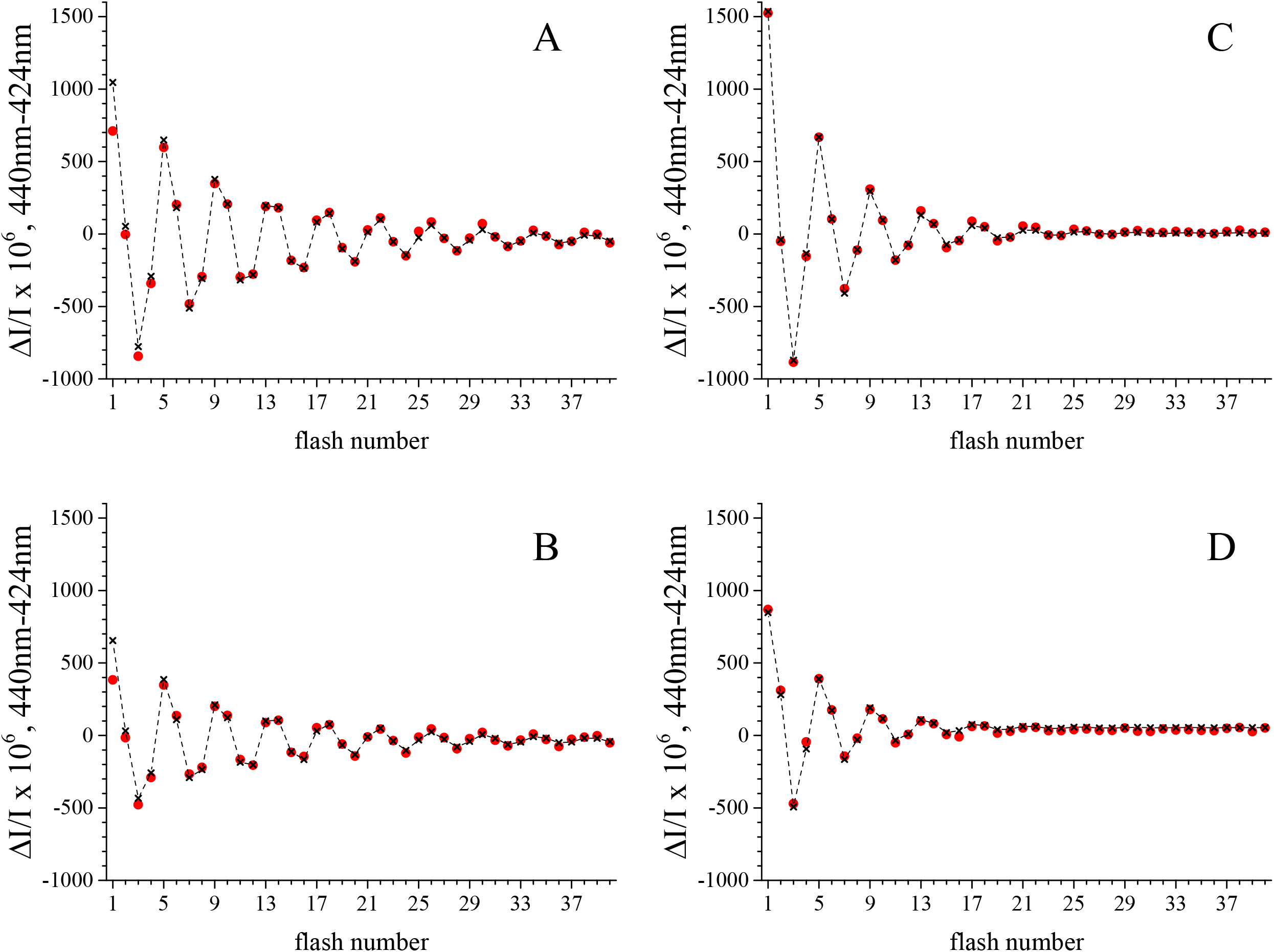
Sequence of the amplitude of the 440 nm-*minus*-424 nm differences using a series of saturating flashes (spaced 400 ms apart). The red full circles, shows the experimental data and the black crosses joined by a continuous line are the fits of the data with the parameters listed in Table 1. Panel A, PSII/Ca at pH 6.0; Panel B, PSII/Sr at pH 6.0; Panel C, PSII/Ca at pH 6.0; Panel D, PSII/Sr at pH 7.0. The samples ([Chl] = 25 µg mL^-1^) were dark-adapted for 1 h at room temperature before the addition of 100 µM PPBQ dissolved in dimethyl sulfoxide. The ΔI/I were measured 100 ms after each flash.

**Figure 4:**
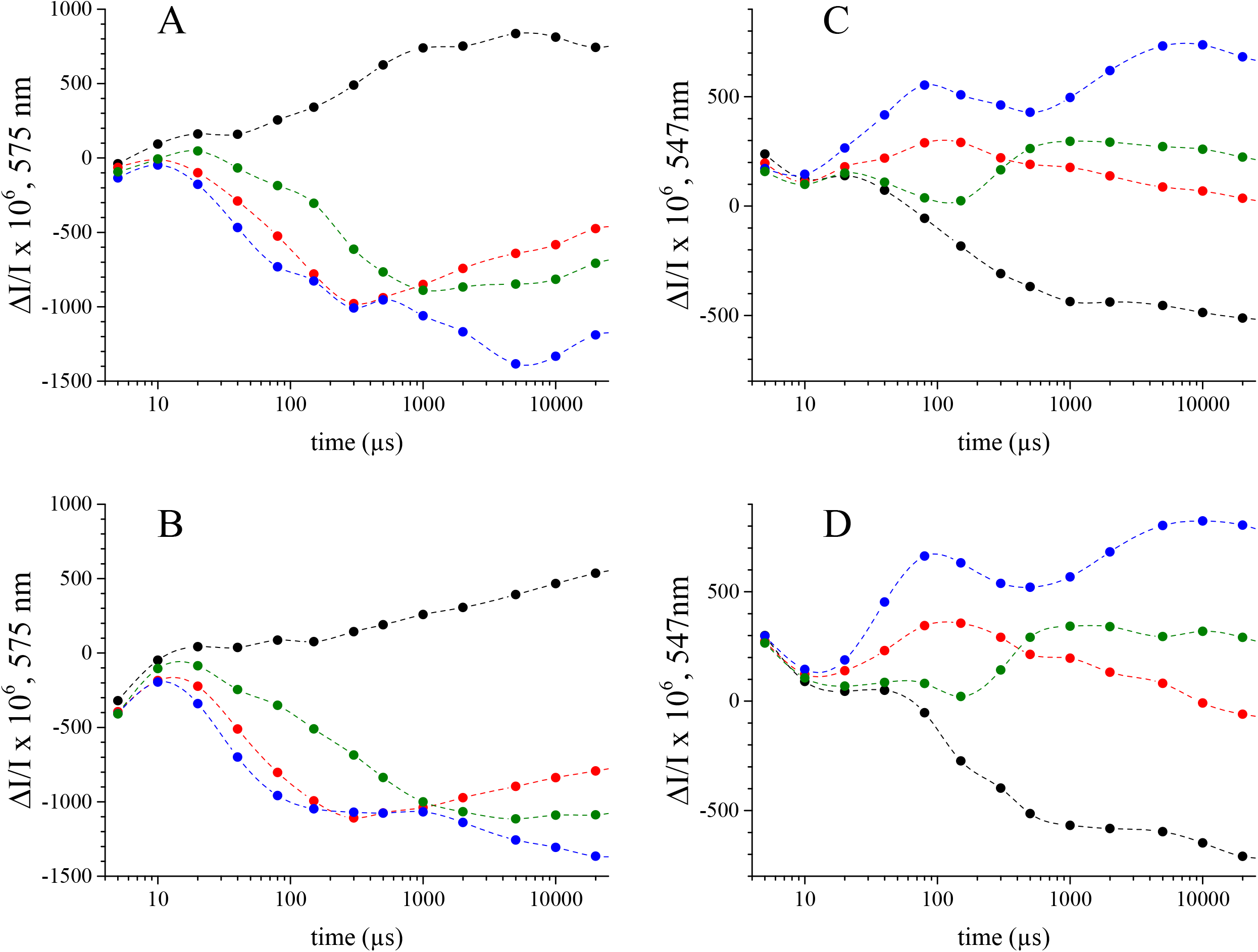
Time-courses of the absorption changes of bromocresol purple at either 575 nm (Panels A and B) or neutral red at 547 nm (Panels C and D) after the 1^st^ flash (black points), the 2^nd^ flash (red points), the 3^rd^ flash (blue points), and the 4^th^ flash (green points) given to either dark-adapted PSII/Ca (Panels A and C) or PSII/Sr (Panels B and D). The dashed lines are spline curves joining the experimental data points. Same other conditions as in Figure 1.

The correspondence between the ΔI/I values calculated in Table 1 in PSII/Ca at pH 7.0 with neutral red and the number of proton released was also estimated by following the OD changes of this dye at 547 nm with a UV-vis spectrometer (UVIKON-XL) between pH 7.04 and pH 6.95 upon the successive addition aliquots of HCl at 38 mM (not shown). With this procedure, we found that the ΔI/I value of 3738 10^-6^ found for a full cycle of the enzyme was equivalent to the addition of 3.5 protons into the bulk which is in very good agreement with the value of 3.6 estimated for a miss parameter of 10 %. Although some simplifications in this control potentially decrease the accuracy of the comparison, it gives significant credit to the quantification of the number of protons released by PSII. It also shows that PSII has little or no buffer capacity. That said, if this had been the case, one could not observe the oscillations with a period of 4 with identical parameters from the beginning to the 40^th^ flash.

### Flash-dependence of the 440 nm-*minus*-424 nm electrochromism

A way to follow the charge(s) in and around the Mn_4_(Ca/Sr)O_5_ cluster is to record the electrochromic band-shifts in the Soret region of the P_D1_ absorption spectrum at 440 nm. This measurement takes into account both the proton uptake/release and the electron transfer events, *e.g.* [60, 62] and references therein. For the removal of the contributions due to the reduction of Q_A_ the ΔI/I at 424 nm was also measured. Indeed, the electrochromism due to Q_A_^-^ equally contributes at 440 nm and 424 nm [72]. The amplitude of the 440 nm-*minus*-424 nm difference was then plotted. Unlike the situation with the dyes, the electrochromism is not contaminated by the reduction/oxidation of the non-heme iron and the coupled protonation/deprotonation [73, 74]. The simulation of the oscillations was therefore done without this contribution as described previously for the oscillations at other wavelengths [63, 66].

Fig. 3 shows the 440 nm-*minus*-424 nm differences induced by each flash of a series given on dark-adapted PSII and measured 100 ms after these flashes. Data in Panels A and B were obtained at pH 6.0 and data in Panels C and D at pH 7.0. The samples were PSII/Ca in Panels A and C and PSII/Sr in Panels B and D. The red full circles are the experimental data and the crosses joined by dashed lines are the results of the fitting procedure. Table 1 summarizes the results from the fittings.

At pH 6.0, the oscillations with a period of four were clearly detectable at least until the 40^th^ flash in both PSII/Ca and PSII/Sr. At pH 7.0, although the oscillations persisted at least until the 40^th^ flash in Fig. 1, the oscillations of the electrochromism significantly decreased after the 25^th^ flash in both PSII/Ca and PSII/Sr. In the four cases, the larger electrochromism is detected on the 1^st^, 5^th^, 9^th^ and so on, flashes, *i.e.* in the S_1_Tyr_Z_^●^ to S_2_Tyr_Z_ transition when the oxidation of the Mn_4_ cluster is not, or only partially, compensated by a proton release. In the four samples, after the 2^nd^ flash and the 4^th^ flash, *i.e.* in the S_2_Tyr_Z_^●^ to S_3_Tyr_Z_ and S_0_Tyr_Z_^●^ to S_1_Tyr_Z_ transitions, the electrochromism is minimum, as expected, since a proton release occurs with the oxidation of the Mn_4_ cluster. After the 3^rd^ flash, *i.e.* in S_3_Tyr_Z_^●^ to S_0_Tyr_Z_ transition, the electrochromism is negative because the cluster is fully reduced and 1 to 2 protons are released. Nevertheless these similarities, small but significant differences are visible, particularly in PSII/Sr at pH 7.0. Indeed, after the first flash, the amplitude of the absorption change was relatively smaller than in PSII/Ca at pH 7.0. In compensation, after the 2^nd^ flash, the change in the absorption was positive in PSII/Sr and negative in PSII/Ca. These differences either did not exist or were less pronounced at pH 6.0 and it is tempting to explain them by a release of proton in a fraction of the centres in PSII/Sr at pH 7.0 after the 1^st^ flash as suggested by the result of the experiment reported in Figs. 1 and 2. These observations are further analysed with the kinetics shown in Fig. 5.

**Figure 5:**
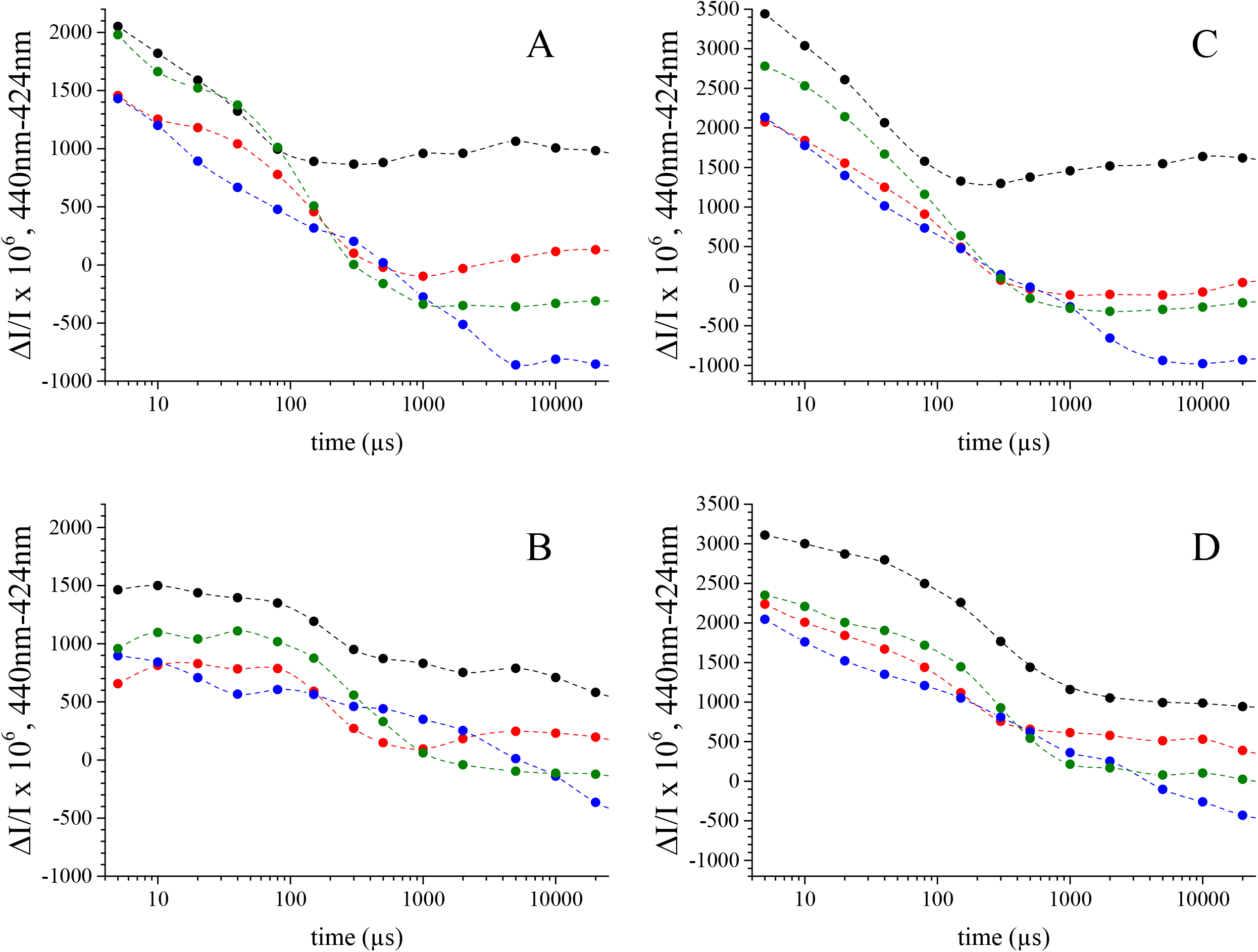
Time-courses of the absorption change differences 440 nm-*minus*-424 nm after the 1^st^ flash (black), the 2^nd^ flash (red), the 3^rd^ flash (blue), and the 4^th^ flash (green) given to either dark-adapted PSII/Ca (Panels A and C) or PSII/Sr (Panels B and D) at either pH 6.0 (Panels A and B) or pH 7.0 (Panels B and D). The dashed lines are a spline curves joining the experimental data points. Same other conditions as in Figure 3.

### Kinetics of the S-state dependent proton release

Fig. 4 shows the kinetics of the decays of the absorption changes of either *i*) bromocresol purple at pH 6.0 with PSII/Ca (Panel A) and PSII/Sr (Panel B) or *ii*) neutral red at pH 7.0 with PSII/Ca (Panel C) and PSII/Sr (Panel D). The measurements were done after the 1^st^ (black points), the 2^nd^ (red points), the 3^rd^ (blue points) and the 4^th^ (green points) flashes given on dark-adapted PSII. The dashed lines joining are spline curves plotted for a better visualisation of the kinetics.

In PSII/Ca, at pH 6.0 (Panel A in Fig. 4), the kinetics were very similar to those already described at pH 6.3 [67]. After the first flash, the increase in the ΔI/I with a *t*_1/2_ close to 300 µs corresponds to the proton uptake following the reduction of the non-heme iron by Q_A_^-^ which occurs with a *t*_1/2_ ∼ 50 µs [10]. After the second flash, a proton release occurred with a *t*_1/2_ close to 60 µs. After the third flash, a biphasic proton release kinetics was resolved. The fastest phase decayed with a *t*_1/2_ of ∼ 40 µs and the slowest one decayed with *t*_1/2_ ≈ 1-2 ms. These two phases in the proton release likely corresponds to the two steps in the S_3_Tyr_Z_^●^ → (S_3_Tyr_Z_^●^)’ → S_0_Tyr_Z_ transitions, *e.g.* [56, 72]. Since the number of protons released in this transition is 1.68 (Table 1) the data can be interpreted with 1 proton released in the first of these two steps and 0.68 in the second one. After the 4^th^ flash, *i.e.* in the S_0_Tyr_Z_^●^ to S_1_Tyr_Z_ transition, a proton release occurred with a *t*_1/2_ of ∼ 200 µs. This proton release in the S_0_Tyr_Z_^●^ to S_1_Tyr_Z_ transition is approximately 4 times slower than the oxidation of the Mn_4_CaO_5_ cluster by Tyr_Z_^●^ [56, 60]. For the longest times, the slow drift discussed above is clearly visible. It does not significantly perturbs the interpretation of the kinetics before 1 ms. For example, in this time range, it is clear that the amplitude of the absorption changes are comparable in the S_2_Tyr_Z_^●^ to S_3_Tyr_Z_, S_0_Tyr_Z_^●^ to S_1_Tyr_Z_ and S_3_Tyr_Z_^●^ → (S_3_Tyr_Z_^●^)’ transitions. However, the drift may slightly decrease the amplitude of the slower phase in the S_3_ to S_0_ transition. The ΔI/I changes observed between 5 µs and 10 µs, were likely due electrostatic effects on the dyes [75], as previously discussed [67].

At pH 6.0 (Panel B), the proton uptake after the first flash in PSII/Sr, with a *t*_1/2_ close to 1 ms, was significantly slower than in PSII/Ca. The proton release in the (S_3_Tyr_Z_^●^)’ → S_0_Tyr_Z_ transition was also slowed down from ∼ 1-2 ms to ∼ 5 ms in agreement with previous results upon the Ca/Sr exchange [66,71]. In the 3 other transitions, S_2_Tyr_Z_^●^ → S_3_Tyr_Z_, S_0_Tyr_Z_^●^ → S_1_Tyr_Z_ and S_3_Tyr_Z_^●^ → (S_3_Tyr_Z_^●^)’, the kinetics of the proton release had similar *t*_1/2_ in PSII/Sr and PSII/Ca.

At pH 7.0, the proton uptake on the first flash (black points) in PSII/Ca occurred with a *t*_1/2_ ≤ 200 µs. This proton uptake occurs over a similar time range as the proton release in the other transitions. For example, after the 3^rd^ flash, i.e in the S_3_Tyr_Z_^●^ → (S_3_Tyr_Z_^●^)’ → S_0_Tyr_Z_ transitions, a proton uptake was clearly detected after the proton release which occurs in the S_3_Tyr_Z_^●^ → (S_3_Tyr_Z_^●^)’and before the proton release occurring in the (S_3_Tyr_Z_^●^)’ → S_0_Tyr_Z_ transition. Despite this complication, the kinetics in PSII/Ca at pH 7.0 appeared similar to those observed in PSII/Ca at pH 6.0. Table 2 shows an estimate of the *t*_1/2_ values after each flash. In PSII/Sr, at pH 7.0, the proton uptake on the first flash had a similar rate as in PSII/Ca at pH 7.0 and pH 6.0. Unfortunately, this proton uptake occurring on the first flash makes difficult the detection of the kinetics corresponding to the release of the ∼ 0.5 proton.

**Table 2.** : Approximate *t*_1/2_ values of the kinetics shown in Figures 4 and 5. Approximate *t*_1/2_ values of the kinetics in Figures 4 and 5 after 4^th^, 1^st^, 2^nd^ and 3^rd^ flashes, *i.e.* in the S_0_ to S_1_, S_1_ to S_2_, S_2_ to S_3_ and S_3_ to S_0_ transitions, respectively, in PSII/Ca and PSII/Sr at pH 6.0 and PH 7.0. The sign in brackets indicates a proton uptake with (+) and a proton release with (-).

| Sample | dye |  |  |  | 440 nm-*minus*-424 nm |  |  |  |
| --- | --- | --- | --- | --- | --- | --- | --- | --- |
| | 4<sup>th</sup> flash<br>$S_0 \rightarrow S_1$ | 1<sup>st</sup> flash<br>$S_1 \rightarrow S_2$ | 2<sup>nd</sup> flash<br>$S_2 \rightarrow S_3$ | 3<sup>rd</sup> flash<br>$S_3 \rightarrow S_0$ | 4<sup>th</sup> flash<br>$S_0 \rightarrow S_1$ | 1<sup>st</sup> flash<br>$S_1 \rightarrow S_2$ | 2<sup>nd</sup> flash<br>$S_2 \rightarrow S_3$ | 3<sup>rd</sup> flash<br>$S_3 \rightarrow S_0$ |
| PSII/Ca<br>pH 6.0 | (-) 200 μs | (+) 300 μs | (-) 60 μs | (-) 40 μs, 1-2 ms | 10-20 μs, 200 μs | 20-30 μs | 10-20 μs, 200 μs | 20 μs, 1 ms |
| PSII/Ca<br>pH 7.0 | (-) 200 μs | (+) <200 μs | (-) 60 μs | (-) 40 μs, 2 ms | 10-20 μs, 200 μs | 20-30 μs | 20 μs, 200 μs | 20 μs, 1 ms |
| PSII/Sr<br>pH 6.0 | (-) 300 μs | (+) 1 ms | (-) 60 μs | (-) 40 μs, 5 ms | 500 μs | 200 μs | 20 μs, 200 μs | 10 μs, 5 ms |
| PSII/Sr<br>pH 7.0 | (-) 200 μs | (+) 200 μs | (-) 60 μs | (-) 40 μs, ≤5 ms | 10-20 μs, 300 μs | 200 μs (20 μs?) | 20 μs, 200 μs | 10-20 μs, 5 ms |

### S state-dependent kinetics of the electrochromism 440 nm-*minus*-424 nm

Fig. 5 shows the kinetics of the 440 nm-*minus*-424 nm differences in the four samples. Panels A and B were measured at pH 6.0 and Panels C and D at pH 7.0. The samples were PSII/Ca in Panels A and C and PSII/Sr in Panels B and D. The absorption change differences 440 nm-*minus*-424 nm are shown after the 1^st^ flash (black), the 2^nd^ flash (red), the 3^rd^ flash (blue), and the 4^th^ flash (green) given to dark-adapted samples. The dashed lines joining the data points are spline curves plotted for a better visualisation of the kinetics. Table 2 shows an estimate of the *t*_1/2_ values after each flash.

After the 1^st^ flash (black points), the 440 nm-*minus*-424 nm difference decayed with a *t*_1/2_ close to 20-30 µs in PSII/Ca at both pH 6.0 (Panel A) and pH 7.0 (Panel C). This decay corresponds to the electron transfer from the Mn_4_CaO_5_ cluster to Tyr_Z_ in the S_1_Tyr_Z_^●^ to S_2_Tyr_Z_ transition since no proton release into the bulk is involved here. In PSII/Sr at pH 6.0 (Panel B), and pH 7.0 (Panel D), the global *t*_1/2_ of the decay after the 1^st^ flash was close to 200 µs. However, we cannot discard an additional fast phase with a *t*_1/2_ close to 20 µs at pH 7.0. The phase with a *t*_1/2_ ∼ 200 µs is almost 10 times slower in PSII/Sr than in PSII/Ca. This difference between PSII/Ca and PSII/Sr is slightly larger than that one previously estimated by following the increase of the absorption of the Mn_4_ cluster in the UV [66]. In both PSII/Ca and PSII/Sr, the amplitude of the decay is approximately twice at pH 7.0 than at pH 6.0. This could suggest that the electrostatic constraint, that is induced by the formation of Tyr_Z_^●^ in the S_1_ state, and that is relaxed during the electron transfer in the S_1_Tyr_Z_^●^ as pH 6.0. to S_2_Tyr_Z_ transition, is larger at pH 7.0 than

After the 2^nd^ flash, two phases were previously identified at pH 6.5 [60]: a fast one with a *t*_1/2_ < 20 µs and a slower one with a *t*_1/2_ between 100 and 200 µs. The slow phase, much slower than the proton release measured with bromocresol purple, has been proposed to correspond to the electron transfer from the Mn_4_ cluster in the S_2_ state to Tyr_Z_^●^ and the fast phase to a proton movement around the Mn_4_ cluster possibly triggered by the formation of the S_2_^HS^ state in the presence of Tyr_Z_^●^. In PSII/Ca at pH 6.0, Panel A in Fig. 5, the decay after the 2^nd^ flash (red points) was qualitatively similar to that one described previously at pH 6.5 [60] with a *t*_1/2_ for the fast phase close to 10-20 µs. In PSII/Ca at pH 7.0 (Panel B), although less evident, the decay seemed also biphasic with *t*_1/2_ values close to 20 µs and 200 µs. In PSII/Sr, the kinetics was similar at pH 6.0 (Panel C) and pH 7.0 (Panel D), with the two *t*_1/2_ values close to 20 µs and 200 µs.

After the 3^rd^ flash (blue points), two phases were detected in all cases. There is a consensus to attribute the fast phase to the S_3_Tyr_Z_^●^ to (S_3_Tyr_Z_^●^)’ + H^+^ transition and the slow one to the (S_3_Tyr_Z_^●^)’ to S_0_Tyr_Z_ + H^+^ transition. In PSII/Ca, the amplitudes of the fast and slow phases were similar at pH 6.0 (Panel A) and pH 7.0 (Panel C) and the two *t*_1/2_, ∼ 20 µs and ∼1 ms, were also similar. In PSII/Sr, the *t*_1/2_ of the slow phase increased to ∼ 5 ms at pH 6.0 (Panel B) and pH 7.0 (Panel D). At pH 7.0, in PSII/Sr, a fast phase with a *t*_1/2_ ∼ 20-30 µs was clearly present. This fast phase seemed to have a smaller amplitude at pH 6.0. The amplitude of the decay of the ΔI/I after the 3^rd^ flash was similar in PSII/Ca and PSII/Sr at pH 7.0. In contrast, at pH 6.0, this amplitude was smaller at pH 6.0 in PSII/Sr than in PSII/Ca.

After the 4^th^ flash, *i.e.* in the S_0_Tyr_Z_^●^ to S_1_Tyr_Z_ transition, in PSII/Ca at pH 6.5, two phases were previously identified and discussed [60]. The fast one with a *t*_1/2_ ∼ 10-20 µs corresponds to the electron transfer and the slow one with a *t*_1/2_ ∼ 200 µs to the proton release. This again confirmed that the electron transfer precedes the proton release in this transition [56, 60]. In Fig. 5, the kinetics after the 4^th^ flash (green points) are in full agreement with these conclusions for PSII/Ca at pH 6.0 and pH 7.0. In PSII/Sr at pH 7.0, the slow phase of the 440 nm-*minus*-424 nm difference with a *t*_1/2_ ∼ 200-300 µs fits well with the kinetics of the proton release in Panel B of Fig. 4. Thus, the electron transfer with a *t*_1/2_ close to 10-20 µs also precedes the proton release in the S_0_Tyr_Z_^●^ to S_1_Tyr_Z_ transition of PSII/Sr at pH 7.0. At pH 6.0, the situation seems to differ in PSII/Sr. Indeed, there is likely only one phase in the 440 nm-*minus*-424 nm difference decay with a *t*_1/2_ ∼ 500 µs. This kinetics is either similar or slightly slower than the proton release occurring with a *t*_1/2_ ∼ 300 µs, which suggests that at pH 6.0, in PSII/Sr, the proton release is time-coupled to (or slightly precedes) the electron transfer in the S_0_Tyr_Z_^●^ to S_1_Tyr_Z_^●^ transition.

## Discussion

The configuration of the S_2_ state capable of progressing to S_3_ remains highly debated in the literature, *e.g.* [21,40–42,49,53,55, 76–82]. Experimentally, in PSII from *T. elongatus*, a S_2_^HS^, *S* ≥ 5/2 spin state, with a *g*value close to 4.8-4.9 could reach the S_3_ ^S=3^ state upon an illumination at temperatures ≤ 200 K while the multiline S_2_^S=1/2^ state could not [36, 37]. In these works, the S_2_^HS^ state was observed at high pH and it was suggested that increasing the pH would mimic the electrostatic influence of Tyr_Z_^•^(His^+^) by displacing the S_2_^LS^ ↔ S_2_^HS^ equilibrium to the right as proposed in a computational work [76]. In this model, at normal pH values, the rapidly released proton in the S_2_ to S_3_ transition before the electron transfer occurs [36,56–60], *i.e.* in the S_2_^LS^Tyr_Z_^●^ (His) state, was proposed to correspond to the formation of the S_2_^HS^ state triggered by Tyr_Z_^•^(His^+^) [36]. In such a model, a change in the flash pattern of the proton release is expected when the flash-induced S_2_-state is high-spin instead of low-spin. This is what we have been looking for in this work. For that, we have followed the proton release in PSII/Ca and PSII/Sr at pH 6.0 and 7.0, knowing that at in PSII/Ca at pH 6.0 and 7.0 and in PSII/Sr at pH 6.0 there is no flash induced S_2_^HS^ state. In contrast, about half of the centers in PSII/Sr exhibit S_2_^HS^ signal at *g*∼ 4.8 at pH 7.0. Given the incredible amount of literature describing different sequence of events in the S_2_ to S_3_ transition, it has become impossible, in a normal article, to discuss all of them and, instead, we have focus our attention on those that we feel are most relevant to the present work. For taking into account some pH dependencies not linked to the formation of a S_2_^HS^ state, the data in PSII/Sr at pH 7.0 were compared to those in PSII/Sr at pH 6.0 but also to those in PSII/Ca at pH 7.0 and pH 6.0.

In PSII/Ca at pH 6.0, the flash pattern for the release into the bulk of the 4 protons is 1.20, 0.0, 1.13, 1.68 for the S_0_→S_1_, S_1_→S_2_, S_2_→S_3_, S_3_→S_0_ transitions, respectively (Table 1). This pattern is comparable, but not identical, to that found in plant PSII at pH 6.0 obtained in a similar experiment that was 1.52, 0.05, 1.0, 1.43 [62]. In [62], the value for the S_3_ to S_0_ transition was computed as the complement to 4 of the sum of the 3 other transitions. The main difference in plant PSII when compared to PSII from *T. elongatus* is a higher value for the S_0_→S_1_ transition compensated by a lower value for the S_3_→S_0_ transition. In the FTIR conditions, at pH 6.0, the pattern was found to be 0.94, 0.28, 1.20, 1.57 in PSII from *T. elongatus* [83]. In plant PSII at pH 7.0, the flash pattern for the proton release into the bulk was 1.10, 0.32, 1.0, 1.58 [62]. In PSII/Ca from *T. elongatus*, at pH 7.0, a significantly different pattern is found here with 0.77, 0.02, 1.13, 2.08. However, in PSII/Ca from *T. elongatus* and from plant, the number of protons released in the S_2_ to S_3_ transition is either very weakly or not affected upon increasing the pH value from 6.0 to 7.0. The other similarities between these two PSII observed upon an increase of the pH from 6.0 to 7.0 is a decrease of the number of proton(s) that are released in the S_0_ to S_1_ transition and an increase of the number of proton(s) that are released in the S_3_ to S_0_ transition. The important difference is the release of 0.32 proton in the S_1_ to S_2_ transition in plant PSII whereas in *T. elongatus* the value remains close to 0. This difference is difficult to rationalize because a pH dependence of the S_1_ to S_2_^LS^ transition between 6.0 and 7.0 in plant PSII was not reported in the literature, *e.g.* [84]. Therefore, it more likely originates from differences in the pK, in S_1_ and S_2_, of some of the groups in the long distance H-bond networks involved in the egress of protons into the lumen upon the oxidation of the Mn_4_CaO_5_ cluster that likely involve the extrinsic subunits, which differ between plant and cyanobacteria, *e.g.* [85, 86]. For that reason, it could be better to compare the pH effects in PSII from only one origin, here *T. elongatus*. In addition, there is no study describing the flash pattern for the proton release in PSII/Sr from plant and the properties of the S_2_^HS^ state are different in plant PSII from an EPR point of view.

In PSII/Sr from *T. elongatus*, at pH 6.0, the flash pattern is 1.35, 0.0, 1.11, 1.54 for the S_0_→S_1_, S_1_→S_2_, S_2_→S_3_, S_3_→S_0_ transitions, respectively. These values are very close to those found earlier, at pH 6.3, in PSII/Sr in which Cl^-^ was replaced with Br^-^. Indeed, the pattern was 1.16, 0.02, 1.19, 1.63 [71]. These values are also very close to those found in PSII/Ca which indicates that either the pK values of the groups involved in the proton release are neither affected by the Ca^2+^/Sr^2+^ exchange nor by the Cl^-^/Br^-^ exchange, or that at pH 6.0 we are far from these pK values. This later possibility would agree with the low (≤ 4.5) and high (≥ 9.5) S-state dependent pK values of 3 of the 4 groups proposed to be involved in the different Sn states that were calculated in [62].

In PSII/Sr at pH 7.0 the number of proton(s) released in the S_1_ to S_2_ transition increased to 0.47. This increase seems mainly compensated by a decreased in the S_0_ to S_1_ transition with 1.02 proton at pH 7.0 instead of 1.35 at pH 6.0 when the reference is the PSII/Sr at pH 6.0. However, increasing the pH in PSII/Ca from 6.0 to 7.0 without favoring the formation of S_2_^HS^ also increases the number of proton released in the S_0_ to S_1_ transition so that the step in which there is a missing proton when the S_2_^HS^ is formed remains to be identified with certainty. It should also be noted that in purified plant PSII the increase in the number of protons released in the S_1_ to S_2_ transition when the pH is increased from 6.0 to 7.0 is not compensated by a decrease in the S_2_ to S_3_ transition but by a decrease in the S_0_ to S_1_ transition [62].

The 440 nm-*minus*-424 nm differences take into account the increase/decrease of the electrostatic environment of P_D1_. In the S-state transitions in which the oxidation of the cluster is preceded/followed by a proton release, the 440 nm-*minus*-424 nm difference is expected to be small. Conversely, the 440 nm-*minus*-424 nm difference is expected to be larger when no proton is released. This is what is observed in PSII/Ca at both pH 6.0 and pH 7.0 and in PSII/Sr at pH 6.0 where no proton are released in the S_1_ to S_2_ transition. For a same proton pattern, the 440 nm-*minus*-424 nm differences are smaller at pH 6.0 in PSII/Sr than in PSII/Ca. This shows that the change in the H-bond network around the Mn_4_ cluster induced by the Ca^2+^/Sr^2+^ exchange have a global consequence on the electrostatic environment of P_D1_ in the S-state cycle.

In PSII/Sr at pH 7.0, in the S_1_ to S_2_ transition, the 440 nm-*minus*-424 nm difference is not smaller than at pH 6.0 while 0.47 protons are released. However, whereas in PSII/Ca the ε_1_ extinction coefficient at pH 7.0 is 1.37 times the value at pH 6.0, in PSII/Sr the ε_1_ extinction coefficient at pH 7.0 is only 1.14 times the value at pH 6.0. This suggests that the electrostatic constraint on P_D1_ differs in the S_2_^HS^ and S_2_^LS^ states and therefore that the H bond network is different in the two spin states of S_2_. Such a result is not surprising since, whatever the structural model for the S_2_^HS^ configuration, the distribution of the charges inside and protons inside and around the cluster differs significantly from those in the S_2_^LS^ configuration, *e.g.* [40,42,76,77,79]. We also know that long-range changes in the Mn_4_CaO_5_ environment can promote the stabilization of the S_2_^HS^ state at the expense of the S_2_^LS^ state as in the V185T mutant [69]. Similarly, a mutation of a Mn_4_CaO_5_ ligand that stabilizes the S_2_^HS^ state, as the D170E mutation, also modifies the H-bond network, *e.g.* [87].

The S-state dependent proton release shows a correlation between the formation of the S_2_^HS^ state and the release of a proton in the S_1_ to S_2_ transition. This correlation does not necessarily imply a causality even if that seems reasonable. Under conditions where the S_2_^LS^ state is the stable S_2_ state, this proton release occurs in the S_2_^LS^Tyr_Z_^●^ state since it is well established that it precedes the electron transfer step [56–60]. In the model proposed here, this proton release would result in the formation of the S_2_^HS^Tyr_Z_^●^ state which would be the competent configuration for the formation of S_3_. Under conditions where the S_2_^HS^ state becomes the stable S_2_ state, the release of the proton is proposed to occur in the S_1_ to S_2_ transition. At this step, we cannot discard the possibility that the S_2_^LS^ state is a transient state is a sequence of events as S_1_Tyr_Z_^●^ → S_2_^LS^Tyr_Z_ → S_2_^HS^Tyr_Z_. Because the exchange of Ca^2+^ with Sr^2+^ stabilizes the S_2_ ^HS^ state, in the context of the model proposed in [39, 40], that would mean that in PSII/Sr the pK of O4 increases and/or that of W1, considered to be the deprotonation site via a channel that includes Asp61 [42,88–90], decreases since the apparent pK of the *g*∼ 2.0 to *g*∼ 4.8 conversion is lower in PSII/Sr than in PSII/Ca [36].

In line with this, we have observed that in a small proportion of PSII/Sr, the S_2_-state produced by an illumination at 180 K gave rise to a *g*∼ 4.3 signal (equivalent to the *g*∼ 4.1 signal in PSII/Ca) that is converted into a *g*∼ 4.8-4.9 signal upon a short warming at ∼ 200 K in the dark [91]. This conversion correlates well with the proton movement proposed in [39, 40] that would be blocked at 180 K and allowed at 200 K. Pushing further the interpretation in the context of the Corry and O’Malley model, the *g*∼ 4.1-4.3 signal would correspond to the *S =* 5/2 S_2_-state with O4 protonated and the *g*∼ 4.8-4.9 signal would correspond to the *S =* 7/2 S_2_-state with O4 deprotonated. In addition, preliminary experiments done in plant PSII under conditions in which the S_2_^HS^ form exhibits the *g*∼ 4.1-4.3 signal suggest that this S_2_^HS^ state is unable to progress to the S_3_ state at 198 K and that, even at pH ∼ 8 (unpublished). In PSII/Ca, upon a Cl^-^/I^-^ exchange, an illumination at 198 K results in the formation of a S_2_^HS^ state with a signal at *g*∼ 4.09 that is unable to progress to S_3_ at low temperatures [91]. Since this chloride is proposed to be involved in the function of the “O4” proton channel [90], the inhibition of the *g*∼ 4.09 state (likely a *S =* 5/2 state) to the *g*∼ 4.8-4.9 state (the *S =* 7/2 state) in PSII/Ca with I^-^ could be due an altered function of the “O4” proton channel making the deprotonation of W1 difficult and thus in the inhibition of the formation of the *S =* 7/2 state in the context of the Corry and O’Malley model [39, 40].

Interestingly, a *g*∼ 4.8 EPR signal with a resolved hyperfine structure was recently detected in plant PSII after the removal of the extrinsic proteins [92] and it would be interesting to know if this state is able to progress to S3 at low temperatures.

In a recent computational work, the consequences of the Ca^2+^/Sr^2+^ exchange on the S_2_ to S_3_ transitions has been analyzed [93]. In this model, two transient states with a closed cubane S_2_^HS^ exhibiting a *g* = 4.1 configuration have been probed. It was found that the more stable S_2_^HS^ state, in PSII/Ca, was with W3 = OH^-^ (with pK = 6.5) and with a neutral His190 whereas in PSII/Sr the more stable state was with W3 = OH_2_ (with pK = 10.3) and with His190(H^+^). If we do not consider the validity or not of the closed cubane option, something that is not a minor issue, these findings, which take into account principally the pK of W3, seems to us difficult to reconcile with our experimental data since in PSII/Ca we would expect W3 as largely deprotonated at pH 8.0 with a pK value of 7 whereas the S_2_ state is mostly in the low-spin configuration [36].

A proton leaving W1 is, in several models, proposed to be transferred to either the Asp61, *e.g.* [40,76,80], or to a close by water molecule (W19 in [40]) thus replacing the H^+^ of W19 which has moved onto O4. In other mechanisms [21, 77] a proton moving towards either Asp61 or W25 would originate from W3, before W3 leaves the Ca^2+^ and binds to Mn1. In all cases, the proton that is released into the bulk cannot be the proton moving onto either Asp61 or W19 or W25, but a proton connected to these proton acceptors via a H-bond network reorganized differently at pH 7.0 in PSII/Sr and PSII/Ca. That would explain why the proton released in the S_1_ to S_2_^HS^ transition in PSII/Sr at pH 7.0 seems not missing in the S_2_ to S_3_^HS^ transition.

The issue concerning the proton and water movements in the S_2_ to S_3_ transition has been addressed recently by following the structural changes in the water and proton channels at times 50 µs, 150 µs, 250 µs, after the second flash [22]. The pieces of information obtained are impressive and impossible to be listed here. One of the conclusions is that the “O1” channel is more likely a water channel than a proton channel. This in agreement with the analysis of the Arg323Glu mutant in which the dielectric relaxation of the protein in the Tyr_Z_ to P_680_^+^ electron transfer is delayed whereas no change in the proton release are observed [67]. It remains that the state, identified in [22], 50 µs after the second flash in the S_2_ to S_3_ transition could correspond to the structural arrangement favoring the formation of the S_2_^HS^ state. Indeed, 50 µs after the second flash, the proton release has already occurred almost completely. Secondly, the structural changes observed at longer times than 50 µs are those accompanying the electron transfer between the Mn_4_CaO_5_ complex and Tyr_Z_.

Table 2 shows the *t*_1/2_ of the kinetics for the proton releases and for the 440 nm-*minus*-424 nm differences from the curves in Figures 4 and 5. The t_1/2_ values are the time at half of the decay of the kinetics determined without a fitting. It should be kept in mind that the kinetics of the 440 nm-*minus*-424 nm differences concerning the proton release steps are necessarily faster than the kinetics of the appearance of the proton into the bulk which is the last step of the proton egress mechanism, see [60, 62].

Table 2 shows that the proton uptake associated with the reduction of the non heme iron after the 1^st^ flash is affected in PSII/Sr at pH 6.0, *i.e.* at a pH value where there is no contamination of the kinetics by a proton release in contrast to the situation at pH 7.0. In PSII/Ca, a pH dependence between 6.0 and 7.0 is not observed. Unfortunately, there is no report in the literature about the electron transfer from Q_A_^-^ to the oxidized non-heme iron in PSII/Sr that could help us to understand the origin of this pH dependence. It was reported that the Ca^2+^/Sr^2^+ exchange slightly up-shifted the *E*_m_(Q_A_/Q_A_^-^) by ∼ +27 mV [94], see however [95]. This increase could slightly slow down the reduction of oxidized non-heme iron at pH 6.0 when compared to PSII/Ca. However, the pH effect observed here on the rate of the proton uptake in PSII/Sr cannot be related to the pK values of the D1-H215 and Glu/Asp residues responsible for the *E*_m_ of the non-heme iron [96]. The positive point remains that at pH 7.0, the kinetics of the proton uptake are similar in PSII/Ca and PSII/Sr, which allows an easier comparison of the kinetics of the proton releases.

After the 3^rd^ flash, the release of the first proton in the S_3_ to S_0_ transition, *i.e.* in the S_3_Tyr_Z_^●^ → (S_3_Tyr_Z_^●^)’ step, occurs at a similar rate in the four samples with a *t*_1/2_ of 40 µs. This *<u>t</u>*_1/2_ is in agreement with the fastest phase in the 440 nm-*minus*-424 nm difference. The kinetics of the 2^nd^ proton in PSII/Sr at pH 7.0, *i.e.* in the (S_3_Tyr_Z_^●^)’ → S_0_Tyr_Z_ step, with *t*_1/2_ ≤ 5 ms, seems faster than expected from the known slowing down from 1-2 ms in PSII/Ca to 5 ms in PSII/Sr for this step [66] whereas this slowing down is observed in the kinetics of the 440 nm-*minus*-424 nm difference at both pH 6.0 and pH 7.0. There is no clear explanation now for that and this observation deserves to be further studied in the future because it is a potential source of information on the mechanism of water oxidation. At least, it is not due to a drift of the pH value. Indeed, i) above pH 7.0 the sample is rapidly damaged something that is not supported by the kinetics after the 1^st^, 2^nd^ and 4^th^ flashes and ii) below pH 7.0 the kinetics of the proton release would be similar to that of the electron transfer something not observed.

In the the S_0_Tyr^●^ to S_1_Tyr_Z_ transition, *i.e.* after the 4^th^ flash, the proton release occurred with a similar rate in the four samples with a *t*_1/2_ of 200-300 µs. In this S-state transition, the proton release in PSII/Ca at pH 6.0 and pH 7.0 and in PSII/Sr at pH 7.0 occurs after the electron transfer that has a *t*_1/2_ of 10-20 µs as previously discussed in detail versus the literature [60]. In PSII/Sr at pH 6.0 the kinetics of the 440 nm-*minus*-424 nm difference seems to have a single phase with a *t*_1/2_ of 500 µs whereas the proton is released with a *t*_1/2_ of 300 µs. This could suggest that at pH 6.0, in PSII/Sr, the proton release either precedes or occurs at the same time as the electron transfer at the opposite of the situation in PSII/Ca and PSII/Sr at pH 7.0.

### Conclusions

Both the stoichiometry and the kinetics of the proton release were followed in PSII/Ca and PSII/Sr at pH 6.0 and pH 7.0. At pH 7.0, the PSII/Sr exhibits a S_2_^HS^ configuration in half of the centers whereas in PSII/Ca at pH 7.0 as at pH 6.0 and in PSII/Sr at pH 6.0 the PSII is in a S_2_^LS^ configuration after 1 flash. Several differences between PSII/Ca and PSII/Sr not yet reported in the literature or better seen here, were found. Among them, the 440 nm-*minus*-424 nm kinetics clearly show that the Ca^2+^/Sr^2+^ exchange is slowing down by a factor 10 the S_1_Tyr_Z_^●^ → S_2_ transition. Previous measurements at 291 nm [66] were less accurate because at this wavelength the absorption coefficient of S_1_Tyr_Z_^●^ and S_2_ are close. In addition, in the S_0_ to S_1_ transition, the electron transfer seems slowed down in PSII/Sr at pH 6.0 to occur almost coupled to the electron transfer in contrast to PSII/Ca at pH 6.0 in which the proton is released after the electron transfer.

The fittings of the oscillations with a period of four strongly indicate that, when half of the centers are in S_2_^HS^ after one flash illumination, about 0.5 proton is released in the S_1_ to S_2_ transition. It is therefore suggested that the proton that is released into the bulk in the S_2_ to S_3_ transition, when the S_2_^LS^ state is the most stable configuration, is released in the S_2_^LS^Tyr_Z_^●^ → S_2_^HS^Tyr_Z_^●^ transition before the electron transfer from the cluster to Tyr_Z_^●^ occurs. Such a model would imply that this proton would be missing in the following S_2_^HS^ to S_3_ transition. Instead, it is principally missing either in the S_3_ to S_0_ transition or in the S_0_ to S_1_ transition while the global charge probed by the electrochromism of P_D1_ increases in the S_2_^HS^ to S_3_ transition in PSII/Sr at pH 7.0 thus suggesting that less proton(s) are released. Further studies are required for a better understanding of the mechanism that links the proton movements in and around the cluster and the proton movements in the channels ultimately resulting by the release of protons into the bulk.

## Acknowledgments

This work has been in part supported by (i) the French Infrastructure for Integrated Structural Biology (FRISBI) ANR-10-INBS-05, (ii) the Labex Dynamo (ANR-11-LABX-0011-01) and (iii) the JSPS-KAKENHI Grant in Scientific Research on Innovative Areas JP17H064351 and a JSPS-KAKENHI Grant 21H02447. This is the last article on the topic of PSII oxygen evolution in my (AB) career. This is an opportunity for me to thank all my coauthors during these 40 years; first and foremost Miwa Sugiura and Bill Rutherford, but also Fabrice Rappaport who passed away too much early, Catherine Berthomieu, Rainer Hienerwadel, Julien Sellés and many other colleagues whose list would be too long here but who, I hope, will recognize themselves.

